# Physiological responses of plants to *in vivo* XRF radiation damage: insights from anatomical, elemental, histochemical, and ultrastructural analyses

**DOI:** 10.1101/2022.01.18.476760

**Authors:** Gabriel Sgarbiero Montanha, João Paulo Rodrigues Marques, Eduardo Santos Rodrigues, Michael W. M. Jones, Hudson Wallace Pereira de Carvalho

## Abstract

X-ray fluorescence spectroscopy (XRF) is a powerful technique for the *in vivo* assessment of plant tissues. However, the potential X-ray exposure damages might affect the structure and elemental composition of living plant tissues leading to artefacts in the recorded data. Herein, we exposed soybean (*Glycine max* (L.) Merrill) leaves to several X-ray doses through a polychromatic benchtop microprobe X-ray fluorescence spectrometer, modulating the photon flux by adjusting either the beam size, focus, or exposure time. The structure, ultrastructure and physiological responses of the irradiated plant tissues were investigated through light and transmission electron microscopy (TEM). Depending on the dose, the X-ray exposure induced decreased K and X-ray scattering intensities, and increased Ca, P, and Mn signals on soybean leaves. Anatomical analysis indicated necrosis of the epidermal and mesophyll cells on the irradiated spots, where TEM images revealed the collapse of cytoplasm and cell-wall breaking. Furthermore, the histochemical analysis detected the production of reactive oxygen species, as well as inhibition of chlorophyll autofluorescence in these areas. Under certain X-ray exposure conditions, *e*.*g*., high photon flux and exposure time, XRF measurements may affect the soybean leaves structures, elemental composition, and cellular ultrastructure, and induce programmed cell death. These results shed light on the characterization of the radiation damage, and thus, help to assess the X-ray radiation limits and strategies for *in vivo* for XRF analysis.

**Highlight:** By exposing soybean leaves to several X-ray doses, we show that the characteristic X-ray induced elemental changes stem from plants’ physiological signalling or responses rather than only sample dehydration.

## INTRODUCTION

Understanding plant functioning is crucial for increasing their efficiency in providing food, fibre, energy, and raw materials for the industry, as well as fostering environment preservation. In this scenario, X-ray fluorescence spectroscopy (XRF) has been widely employed to explore the elemental composition and spatial distribution in a myriad of botanical materials (van der Ent *et al*. 2018; Gomes *et al*. 2019; Montanha *et al*. 2020a; Lau *et al*. 2020; Romeu *et al*. 2021; Macedo *et al*. 2021). New generations of XRF instrumentation, *i*.*e*., faster high energy-resolution detectors (van der Ent *et al*. 2018; Montanha *et al*. 2020b) and portable XRF systems enabled to assess plant specimens under *in vivo* and *situ* conditions (Montanha *et al*. 2020b; Montanha *et al*. 2020c; Mijovilovich *et al*. 2020; Soares *et al*. 2021; Corrêa *et al*. 2021), thereby providing means for tracking ever-changing physiological processes, *e*.*g*., the absorption of nutrients and pollutants through roots and leaves, and their movement toward tissues while they are taking place (van der Ent *et al*. 2018; da Cruz *et al*. 2019; Montanha *et al*. 2020b; Montanha *et al*. 2020c).

XRF is often regarded as a non-destructive tool. Nevertheless, X-ray beam-induced radiation damage is a critical issue for the *in vivo* X-ray analysis of biological samples. In principle, the X-ray ionizing feature can affect either the composition or structure of the materials, thereby producing artefacts that might result in biased results (Smith *et al*. 2009; Matsushima *et al*. 2013; Terzano *et al*. 2013). X-ray radiation can lead to cell damage, decrease the photosynthetic activity (Matsushima *et al*. 2013), photoreduction of elements (Smith *et al*. 2009), changes in element distribution and breaking of chemical bonds, thus, also altering the chemical speciation (Terzano *et al*. 2013). The type and extent of radiation damage mainly depend on the inherent characteristics of the samples, *e*.*g*., chemical composition and state, and the analysis conditions such as the beam energy, flux density, and the measurement’s dwell time (Vijayan *et al*. 2015).

The existence of radiation damage is a well-documented issue during XRF analysis of plants, especially on those carried out on synchrotron sources (Ellis *et al*. 1998; Freeman 2006; A. Castillo-Michel *et al*. 2016; Jones *et al*. 2017; Gomes *et al*. 2019). On the other hand, it also poses an issue with laboratory devices (van der Ent *et al*. 2018; Jones *et al*. 2019), *e*.*g*., X-ray radiation-induced damage was found on *Arabidopsis thaliana* leaves subjected to *in vivo* XRF mapping in a house-made µ-XRF spectrometer furnished with Rh anode (50 W, 50 kV 1 mA) and focused X-ray beam of 23.7 µm FWHM at WL_3_ edge (Fittschen *et al*. 2017). These studies focused on characterising the changes in the elemental composition as a function of X-ray exposure. However, it is still unclear whether the X-ray induced stress also leads to plants’ physiological responses.

In this scenario, the present study systematically investigated the effects of X-ray irradiation on soybean (*Glycine max* (L.) Merrill) leaves. Besides monitoring the redistribution of elements during the X-ray irradiation, the effects on tissue morphology and biochemistry were also investigated to provide a comprehensive overview of the plant’s response to XRF analyses.

## MATERIALS AND METHODS

### Plant materials and cultivation

The experiments were carried out using soybean plants (*Glycine max* (L.) Merrill) at the V3 growth stage, *i*.*e*., when the second trifoliate leaf was completely expanded. The soy seeds (M7739 IPRO variety, Monsoy, Brazil) were sowed in plastic pots containing a moistened sandy substrate and irrigated with deionised water during the first seven days, and thereupon, with 1:5 Arnon’s and Hoagland hydroponic solution (Hoagland and Arnon 1950) until the plants reached the V3 growth stage. The cultivation took place in a growth room under a 12 h-photoperiod at 250 μmol of photons m^-2^ s^-1^ and 27 ± 2 °C. Each experiment herein described was carried out using at least two independent biological replicates.

### In vivo X-ray radiation exposure assays

The irradiation assays were conducted on the cotyledonary leaves. The soybean plants were transferred to an acrylic sample holder conserving the substrate, with the leaves fixed with Kapton tape on the top of a 5 µm polypropylene film (Spex SamplePrep, no. 3520, USA) and loaded into a benchtop microprobe X-ray fluorescence equipment (µ-XRF, Orbis PC EDAX, USA). Supplementary data Fig. S1 details the setup assembled for conducting the experiments. The µ-XRF equipment is furnished with a 50 W Rh X-ray tube at 17.44 keV Mo-Kα energy yielding either a 1 mm collimated X-ray beam (hereby referred to as “low photon flux” condition), or a 30 µm polycapillary focused beam (hereby referred to as “high photon flux”Bcondition. Spectra were recorded at 2 to 4 min intervals for longer studies, and 3 s intervals for the shorter studies. Additional tests were conducted using a 25-µm thick Ti primary filter to enable Mn detection. The spectra were recorded by a 30 mm^2^ silicon drift detector with a deadtime below 10%. The elemental signals were distinguished from the background using their threshold values (*T*), calculated according to equation 1;

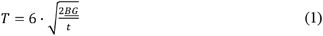

where <u>BG</u> (cps) is the average of 10 random measurements of background count rate under the corresponding analyte signal, and *t* (s) is the acquisition time (Jenkins 1995; Kadachi and Al-Eshaikh 2012).

The X-ray doses deposited into the samples (Gy) were calculated according to (Jones *et al*. 2017);

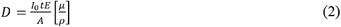

Where: I_0_ is incident photon intensity (ph s^-1^), t is the measurement time (s), *E* is the weighted average energy of X-ray spectra at each instrumental condition employed in the study, A is sampling area (m^2^) and 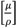 is the X-ray mass attenuation coefficient of the soybean leaf sample (m^2^ kg^-1^)

Since the study explores a polychromatic X-ray source, the doses were estimated by using the weighted average energy of the X-ray spectra for each instrumental condition employed in the study irradiating a plexiglass cube with and without a 100 µm thick Rh primary filter.

The incident X-ray beam fluxes (*I*_*0*_), in photons s^-1^, were determined at the Rh Kα emission line using an electrometer system (Keithley, model 6514, USA) coupled with a 10 µm-thick Si photodiode (Alibava, Spain), calculated according to (Jones *et al*. 2017);

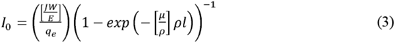

where J is current flowing through the photodiode detector (C s^-1^), the work function of the detector material, *i*.*e*., 3.6 eV, *E* is the Rh Kα emission line photon energy, *i*.*e*., 20216 eV, *q*_*e*_ is the elementary charge, *i*.*e*., 1.60×10^−19^ C, *l* is the photodiode detector active length, *i*.*e*., 10^−3^ cm, *ρ* is the density of the detector material, *i*.*e*., 2.33 g cm^-3^, and 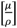 is the mass attenuation coefficient of the detector, *i*.*e*., 3.866 cm^2^ g^-1^.

### XRF elemental characterization of the irradiated plant tissues

Fresh and dried samples were irradiated for 24 min with a high photon flux before being left at rest for 24 hours. The leaf segments were then mapped under *vivo* conditions, detached and dried in a laboratory oven at 60 ºC for 48-h, and reanalysed under the same conditions. The irradiated areas were mapped within a 32 × 25-pixel matrix and scanned along a 32-point line using the benchtop µ-XRF instrument. Both analyses were carried out by employing a 30 µm X-ray beam generated by an Rh X-ray tube at 40 kV and 500 µA, with a 25 µm-thin Ti primary filter selected. The dwell time was 3 s for the maps and 10 s for the linescans, respectively.

### Histochemical characterisation of irradiated plant tissues

To verify whether radiation damage induces physiological changes in the irradiated area, soybean leaves were irradiated for 3 or 240 s with high photon flux. Following irradiation, one group of leaves were detached from the petiole 2 or 24 h past the irradiation and imaged using a digital microscope (Hirox, KH-8700, Japan), whereas another group was subjected to histochemical assays to verify the occurrence of H_2_O_2_ through immersion in a 3,3’-diaminobenzidine (DAB) staining solution (1 mg mL^-1^ DAB, Tween 20 (0.05% v/v) and 10 mM Na2HPO4, pH > 3-4] for 2-4 h, according to the method previously described (Daudi and O’Brien 2012); callose (1,3-β-Glucan) by staining with an aniline blue (0.01% v/v in phosphate buffer, pH 7.2) solution (Brandizzi 2000) for 20 min; and autophagic vacuoles through staining using a monododecyl cadaverine (MDC, 0.001 mol L^-1^ in phosphate buffer) solution for 30 minutes (Contento *et al*. 2005; Kabbage *et al*. 2013). In all tests, the leaf samples were directly mounted on the glass slides containing PBS buffer and analysed through an autofluorescence light microscope (Zeiss, Axion Observer, Germany). The fluorescent images were taken using a filter set (excitation: 365 nm; emission: 420 nm)., which was also employed to detect chlorophyll autofluorescence (Donaldson 2020).

### *Anatomical* characterisation of irradiated plant tissues

A *ca*. 3 mm^2^ piece of the leaf samples irradiated for 240 s at high photon flux were harvested 24 after the exposure, then fixed in Karnovsky’s solution (Karnovsky 1965), dehydrated in an increasing ethanolic series, embedded in historesin, sequentially cross-sectioned in a rotatory microtome, and stained with toluidine blue (TBO, Sigma-Aldrich, Germany), according to the method described elsewhere (Marques *et al*. 2015). The samples were then mounted on glass slides containing synthetic resin (Entelan^®^, Sigma-Aldrich, Germany). The images were taken using a microscope (Zeiss, Axion Observer, Germany).

### Transmission electron microscopy structural characterization of irradiated plant tissues

Following X-ray exposure, the leaves irradiated for 24 min and 10 s with high photon flux were harvested and immediately immersed in a Karnovsky’s fixing solution (Karnovsky 1965), then post-fixed in 1% v/v osmium tetroxide + 0.1 mol L^-1^ cacodylate buffer (pH 7.2). The samples were washed thrice in a cacodylate buffer and transferred to a 0.5% v/v uranyl acetate solution overnight. Thereafter, the leaf samples were washed thrice in water and dehydrated in ketone series (30%, 50%, 70%, 90%, and 100%, twice) for 15 minutes (Marques and Soares 2021) and embedded in Spurr resin (Sigma-Aldrich, Germany) with the standard formulae. The blocks were then cut in an ultramicrotome (UC6, Leica, Germany) and the ultrathin sections were stained with 5 % v/v uranyl acetate and lead 2% v/v citrate for 15 min each (Reynolds 1963). The samples were analysed under transmission electron microscopy (TEM 1011, JEOL, Japan). The elemental, histochemical, anatomical, and ultrastructural characterization of irradiated leaf tissues are summarized in Fig. 1.

**Fig. 1.**
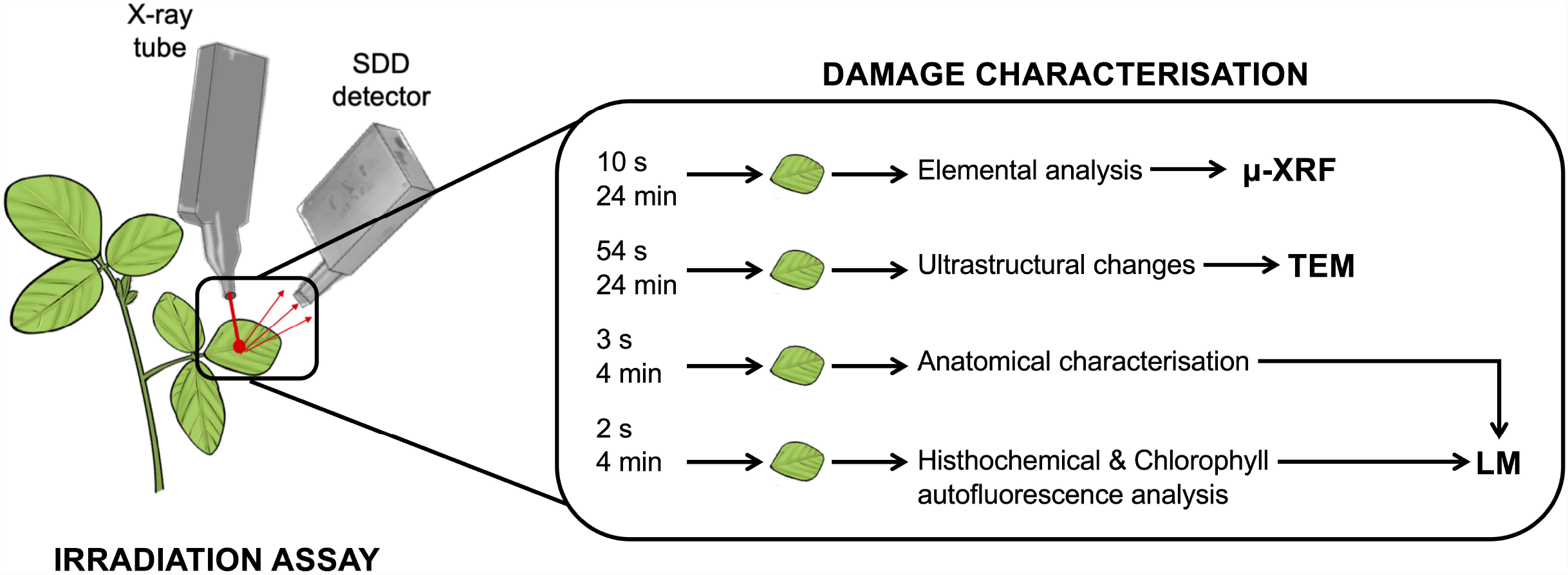
Scheme of the experimental workflow for the elemental, histochemical, anatomical, and ultrastructural characterization of irradiated soybean leaf tissues. Samples irradiated for 2 s and 240 s were subjected to histochemical and anatomical tests and analysed with light microscopy (LM) while samples irradiated for 10 s and 24 min were subjected to ultrastructural tests with TEM and u-XRF.

## RESULTS

### Elemental changes in irradiated soybean tissues

Figure 2A-D shows the XRF signals for the K-alpha peaks of P, K, and Ca, as well as the Rh Lα scattering recorded in living soybean leaves exposed to four different X-ray conditions. These results reveal that the low photon flux (Fig. 2A) does not alter the intensities of the detected elements. However, one should keep in mind that a high photon flux beam is used for generating elemental maps. Under these conditions, one can notice that the effects of radiation damage are significant over 24 min (Fig. 2B), with the onset of damage occurring in less than a minute (Fig. 2C). The general trend is a decrease in K and an increase in Ca, suggesting that the K is being removed from the damaged region while Ca is being accumulated. Reducing the power by an order of magnitude to 4.5 W (Fig. 2D) shows a similar trend as those observed in the exposures at 40.5 W, albeit at a slightly slower rate. In all cases, the Rh Lα scattering did not present any clear pattern of change, suggesting the damage observed is not mass loss (Jones *et al*. 2019). Employing a Ti filter enabled the detection of Mn, showing a sharp increase as a function of the exposure time (Fig. 2E). Supplementary data Fig. S2 presents the same trend obtained on independent biological replicates. In addition, Supplementary data Fig. S3 shows elemental composition modification induced by 24 min irradiation under the 40.5 W focused beam was not reversible after two hours of plant rest. In other words, the tissue was not able to recover itself during this time interval.

**Fig. 2.**
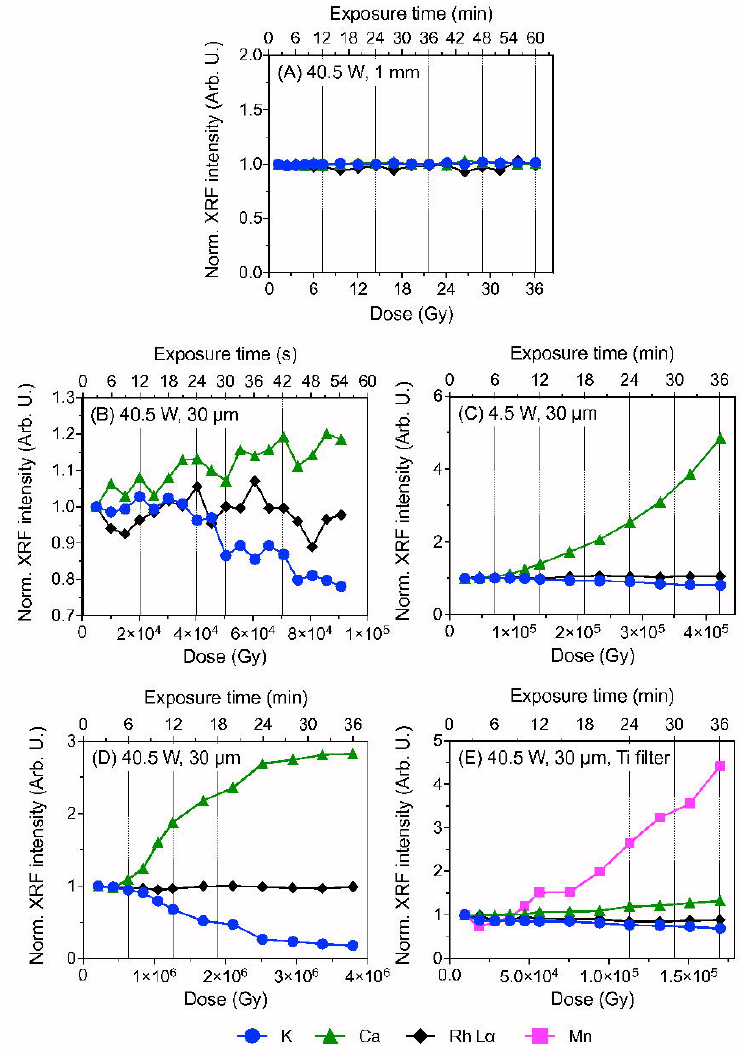
Normalized XRF K, Ca, and Rh Lα count rate recorded on soybean leaves at the V3 growth stage as a function of the time and radiation dose during the exposure either to a collimated 1 mm or polycapillary focused 30 µm X-ray beam derived from an Rh anode at 40.5 W (45 kV and 900 µA, A-C) or 4.5 power (45 kV and 100 µA, D), without (A-D) or with a 25-µm thick Ti primary filter selected (E). The count rates were normalized by their respectively first recorded values.

To distinguish the modifications induced by the X-ray beam from those caused by dehydration, the elemental composition of fresh and 48h oven-dried soy leaves irradiated with the high photon flux were imaged by µ-XRF, presented in Fig. 3. The photographs of the soybean leaf tissues in both cases reveal visual signs of necrosis (dark spot) in the irradiated region 24-h after the X-ray exposure. The chemical maps, in line with the time-resolved exposure (Fig. 2), shows a sharp decrease in K and an increased Ca intensity in the irradiated spots. Similarly, the Mn count rate is *ca*.10 to 50-fold higher in the irradiated regions compared to its neighbouring regions. This is highlighted in the linescans, which provide a higher lateral resolution than the maps. The same trend is observed on the 48-h oven-dried leaves (Fig. 3B), thus, under the available lateral resolution, the tissue drying did not influence the elemental spatial distribution. Therefore, we do not suspect that radiation-induced dehydration is responsible for the trends observed. The same trend was recorded on independent biological replicates presented in Supplementary data Fig. S4. On the other hand, no damage was observed for a 10 s exposure, as expected from data presented in Fig. 2D and Supplementary data Fig. S5.

**Fig. 3.**
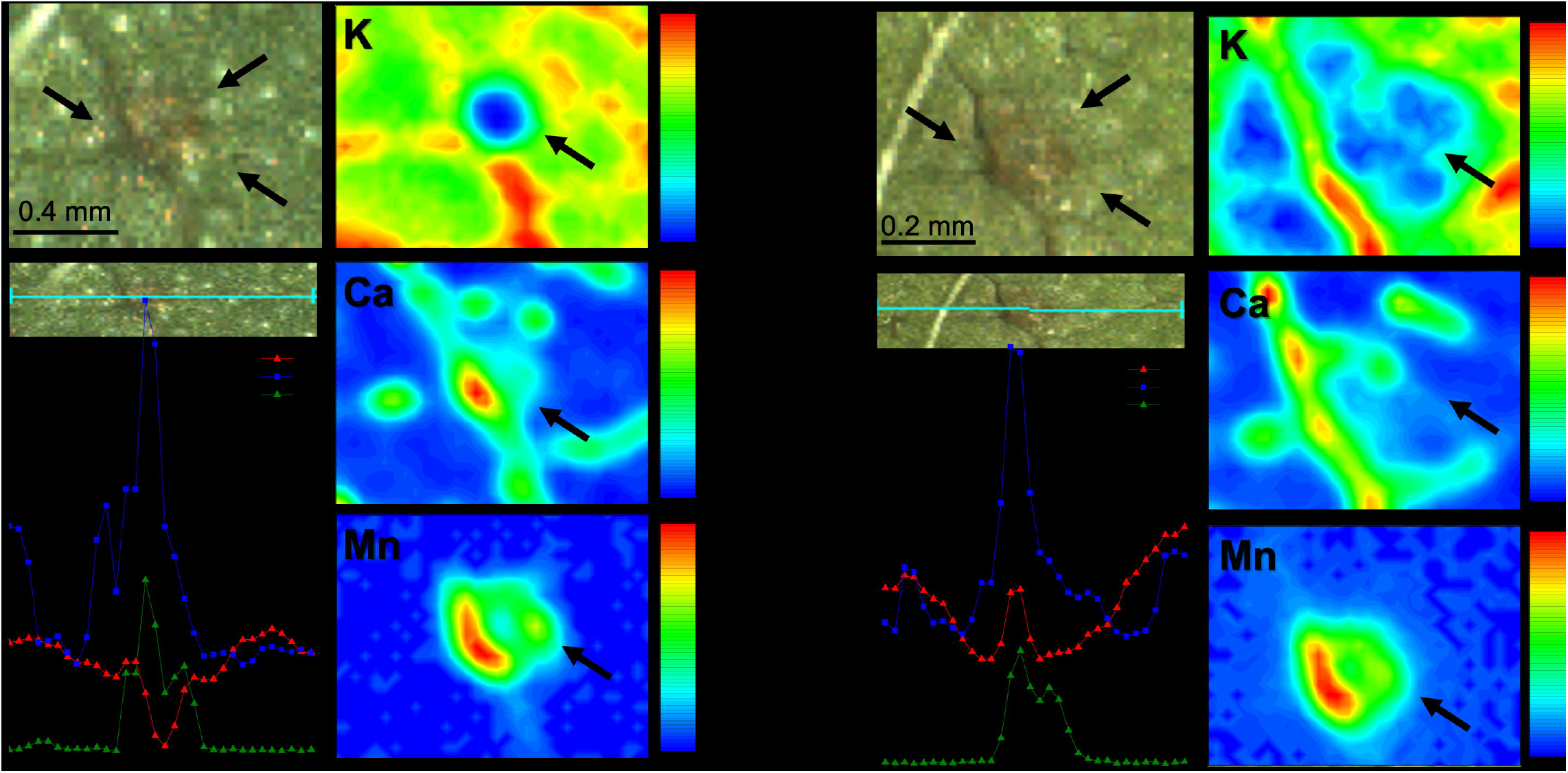
Photographs and XRF probing of K, Ca, and Mn spatial distribution on fresh (A) and dried (B) soybean leaves at the V3 growth stage exposed to polycapillary focused 30 µm X-ray beam at 40.5 W (45 kV and 900 µA) for 24 minutes (2×10^6^ Gy dose). An 800-pixel XRF map and a 32-point XRF linescan were carried out in each case. The fresh leaves were monitored 24 h past the irradiation, then oven-dried at 60ºC for 48 h and reanalysed at the same instrumental conditions. The black arrows denote the irradiated regions, where a necrotic spot and clear changes in the elemental distribution in both fresh and dried leaf tissues are observed.

### X-ray induced *anatomical, histochemical and* ultrastructural changes in soybean leaf tissues

A series of plant anatomy techniques, *i*.*e*., anatomical, ultrastructural, and histochemical, assays were herein explored to properly determine which factors have taken a role in the observed X-ray induced damage presented in Fig. 2 and Fig. 3.

Comparing the anatomical patterns of the irradiated leaf tissues, *i*.*e*., exposed to 10 s (Figs. 4A-F) or 20 min (Figs. 4G-O) to the high photon flux beam, and the non-irradiated tissues, one cannot observe any damage in those regions irradiated for 10 s. These leaflets present a single lens-shaped epidermal cell layer with a thin cuticle (Figs. 4A-E), as well as a dorsiventral mesophyll with 1-2 palisade parenchyma cells and 4 to 5 layers of spongy parenchyma (Fig. 4B) cells with a high number of chloroplasts with starch (Fig 4F). The vascular bundle is collateral and presents a leaf sheath extension (Fig. 4A).

**Fig. 4.**
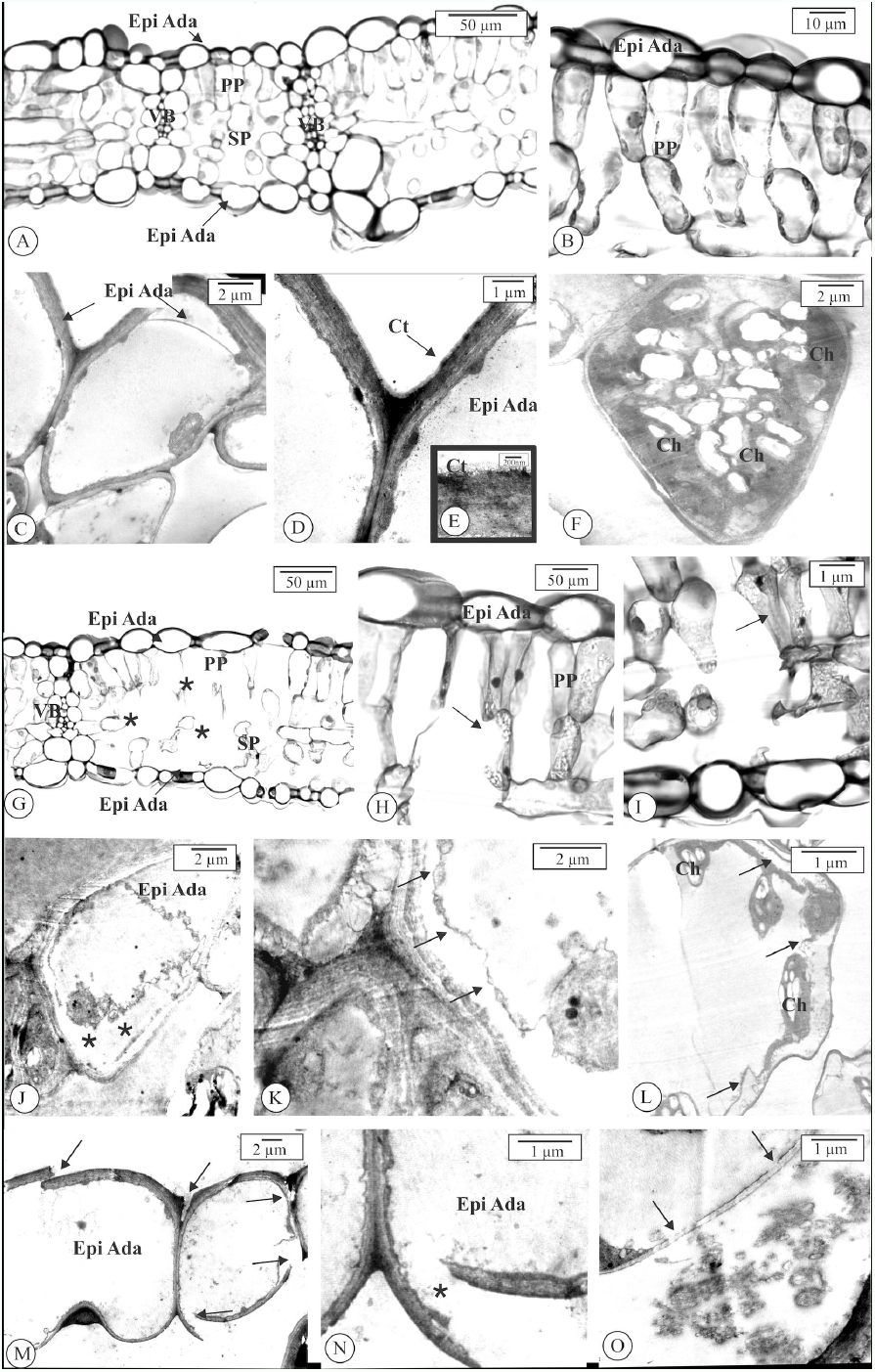
Light and electron micrographs of regions of soybean leaflets at the V4 growth stage irradiated for 10 s (A-F) and 20 min (G-O) with 30 µm focus generated by a Rh anode with 40.5 W of power (45 kV and 900 μA). The anatomy of the 10 s (2×10^4^ Gy) irradiated tissues remains healthy, indicating that epidermal cells are lens-shaped and covered by a thin cuticle (A-E). F. Spongy parenchyma cell with chloroplasts. The 24 min (2×10^6^ Gy) irradiated tissues (G-O) show injured regions (* in G). The lesion promotes cell wall rupture (arrows H; I; M-N). Cell plasmolysis of the leaf and plasm membrane detachment were observed (* in J and arrows in K-L). M. Protoplast leaking in the intercellular space. Ch – chloroplast; Ct – cuticle; Epi Ada – epidermis adaxial; Pp – palisade parenchyma; Sp – spongy parenchyma; Vb – vascular bundle.

Conversely, the damage is observed in the epidermis and mesophyll cells of the leaves irradiated for 20 min. The damaged region encompasses a *ca*. 100 µm area, more than three-fold larger than the putative 30 µm X-ray beam spot size (Fig. 4G). The TEM images revealed the rupture of cell walls in the mesophyll and epidermal cells (Fig. 4H-I; L-N), as well as plasmolysis, *i*.*e*., the detachment of the plasm membrane (Fig. 4J-K), and the leakage of the cytoplasm to the extracellular region (Fig. 4O).

Furthermore, the histochemical tests conducted on soybean leaves either 2 or 24 hours after the exposure (h.a.e) to the high photon flux beam during 2 or 240 s (Fig. 5) unveiled that regardless of the recovery time, the 2 s X-ray shots neither impaired the chlorophyll autofluorescence nor induced detectable production of H_2_O_2_, callose, or autophagic vacuoles (Fig. 5 A-D; I-L), whereas extinction of chlorophyll autofluorescence (Fig. 5 E; M), as well as a positive reaction with both aniline blue and monododecyl cadaverine reagents, were recorded for the 240 s shots, depicting the development of callose and the autophagic vacuoles (Fig. 5G; O and Fig. 5H-P). Oddly, although the positive diaminobenzidine (DAB) reaction confirms the H_2_O_2_ accumulation in the high photon flux irradiated area 2 h.a.e (Fig. 5F-F1), it was not detected 24 h.a.e. (Fig. 5N) suggesting that the induced damages hampered the cells redox metabolism, thus, their ability in producing ROS.

**Fig. 5.**
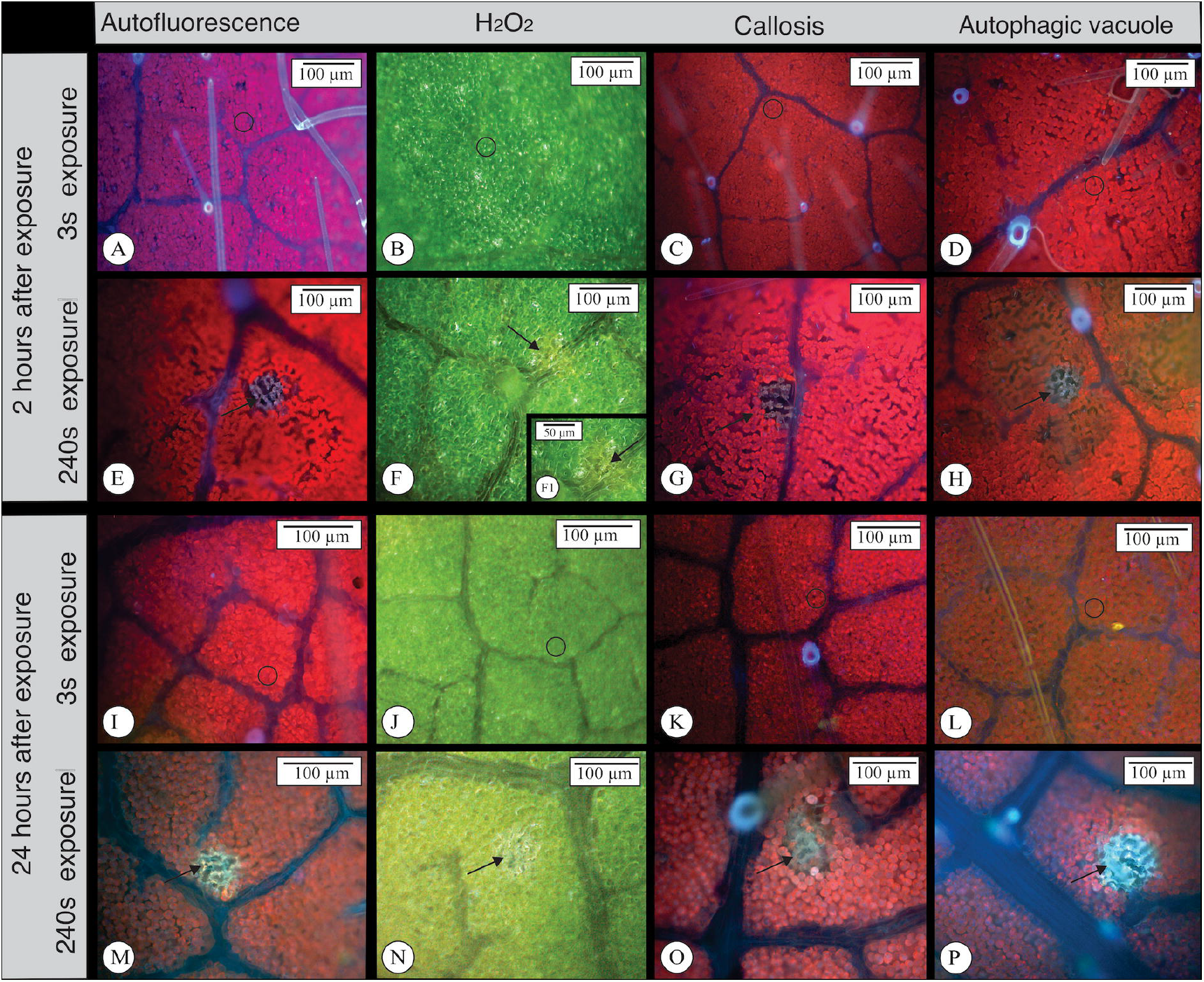
Light microscopy of soybean leaflets exposed to different hours after exposure (h.a.e.) and doses of X-ray and analysed after different histochemical essays. Autofluorescence analysis was also conducted to verify chlorophyll emission extinction. Low exposure (3 s, 5×10^3^ Gy) does not cause any damage to the tissue or induce H_2_O_2_, callose or autophagic vacuole, whereas a high X-ray dose (240 seconds, 4×10^5^) causes a lesion in the leaflet (E-H; M-P) on both, 2 and 24 hours after exposure.

## DISCUSSION

One of the main analytical advantages of X-ray fluorescence spectroscopy is the ability to interrogate the elemental composition and distribution of biological materials in their steady states (Montanha *et al*. 2020b; Montanha *et al*. 2020c; Pushie *et al*. 2020), with minimal or even no sample preparation requirements (Rodrigues *et al*. 2018; Montanha *et al*. 2020b). From a botanical standpoint, the *in vivo* and *situ* XRF-based analysis of plant tissues encompass a unique opportunity to elucidate key mechanisms of plants physiology, such as root-to-shot and foliar nutrient uptake (Gomes *et al*. 2019; Montanha *et al*. 2020b; Montanha *et al*. 2020c; Corrêa *et al*. 2021), metal hyperaccumulation (Hernandez-Viezcas *et al*. 2013; van der Ent *et al*. 2018) and toxicity (Reis *et al*. 2020; Lanza *et al*. 2021), and changes in elemental distribution in healthy and injured tissues (Marques *et al*. 2020; Macedo *et al*. 2021; Naim *et al*. 2021).

Nonetheless, the X-ray beam-induced radiation damage is a major issue for the *in vivo* XRF measurements of biological materials, since the ionising nature of the X-rays can alter both the composition and structure of the samples, producing artefacts that might severely impact the driven results (Smith *et al*. 2009; Matsushima *et al*. 2013; Terzano *et al*. 2013). X-ray induced damage mainly depends on the dose, *i*.*e*., the amount of radiation deposited in a certain volume, which is a function of the incident photon flux intensity, samples’ X-ray mass attenuation coefficient, and the exposure time (Oger *et al*. 2008; Jones *et al*. 2017, 2019). This is highlighted in our irradiation experiments (Fig. 2), which clearly showed that the higher the dose, the higher the changes in the elemental signals of an irradiated spot. Several studies thus far have linked that radiation doses of 10^7^ Gy led to photoreduction and subsequent damage of biological materials, such as metalloproteins (da Cruz *et al*. 2017; Yano *et al*. 2005; Smolentsev *et al*. 2009; James *et al*. 2016). Conversely, our data reveal the onset of elemental redistribution at 10^4^ Gy for fresh soybean leaves. These results are similar to those found by Jones *et al* 2019, which indicated that a 10^2^ Gy might damage sunflower (*Helianthus annuus*) leaves. Together, these results suggest that the radiation limits are usually lower for plant tissues compared to animal ones and that different plant tissues may have different radiation limits.

A sharp decrease of K and an increase of Ca signals were detected whenever employing the polycapillary 30 µm beam (Fig. 2B-E). These findings are quite similar to those results reported in our previous studies (Jones *et al*. 2019; Montanha *et al*. 2020c). Furthermore, the 25 µm-thick Ti primary filter enabled us to record a remarkable increase of Mn signals due to the reduction of the background radiation reaching the X-ray detector. However, the use of this filter between the X-ray source and the sample also reduces the photon flux, thereby leading to less extensive changes in the elemental composition for a given exposure time. This phenomenon is illustrated in Fig. S6, which presents the XRF recorded on a Plexiglas cube with and without the 25 µm-thick Ti primary filter selected. We note that due to the reduced flux, elemental sensitivity is also reduced, and longer exposure times would be required to achieve the same signal as without the filter for elements apart from Mn.

Although it seems that plant materials are prone to tissue dehydration caused by the heating during beam exposure either at a synchrotron facility or laboratory-based XRF equipment, directly interfering with elemental distribution and chemical speciation (Tylko *et al*. 2007; Lombi *et al*. 2011), it is important to highlight that the Rh-Lα scattering peaks, highly sensitive to the samples total mass and therefore dehydration (Bastos *et al*. 2012), was only slightly affected by the X-ray exposure in our assays (Fig. 1). Therefore, dehydration might not be the only, or major, contributing factor to the elemental changes recorded. The elemental maps of the irradiated areas of fresh and dried tissues (Fig. 3) reinforce this hypothesis, since both K, Ca, and Mn distribution pattern remains after sample drying.

Additionally, since the observed changes were not reversible (Fig. 3, Supplementary data Fig. S2-S3), they might likely be related to plant response to the acute stress induced by the X-ray exposure. For example, increased Ca and Mn contents have been observed on wheat leaves inoculated with a fungal pathogen (Naim *et al*. 2021).

The anatomical and ultrastructural analyses revealed that the damaged area encompasses a spot *ca*. 3-fold higher than the 30 µm beam at Mo K-alpha energy (Fig. 4). Although the incident beam herein explored was not a monochromatic one, longer wavelengths result in larger beams. However, this does not account for the extent of the damaged region. Moreover, the histochemical assay with DAB indicated the presence of H_2_O_2_ (Fig. 5B; F; J; N), a key reactive oxygen species (ROS) in plant physiology (Quan *et al*. 2008), 2 h past the 20 min irradiation with the high photon flux, unveiling a localised molecular response to a stressful condition. Interestingly, one should notice that Mn is a cofactor of the Mn-superoxide dismutase, a crucial enzyme in the redox metabolism of plants (Morgan *et al*. 2008), whereas Ca is a well-known marker of abiotic stress in plants (White and Broadley 2003; Toyota *et al*. 2018). It does suggest that the increasing Ca and Mn signals recorded during the irradiation might reflect both stress-like signalling and demand for enzymes for ROS scavenging processes. Moreover, although the presence of callose (β-1,3-glucan), is a cell-wall polysaccharide strongly related to biotic and abiotic stress signalling (Chen and Kim 2009; Zavaliev *et al*. 2011), was not observed in the stress. Moreover, autophagic vacuoles are a specialised group of organelles holding hydrolytic-like enzymes related to cell digestion which plays crucial roles in programmed cell death (PCD) processes (Floyd *et al*. 2015) was also not verified. This suggests that the 20 min X-ray exposure was able to destroy soybean leaf tissues even before callose outbreak and autophagic vacuoles production, as emphasized by the cell wall ruptures, plasmolysis and protoplast leaking (Fig. 4 and 6).

**Fig. 6.**
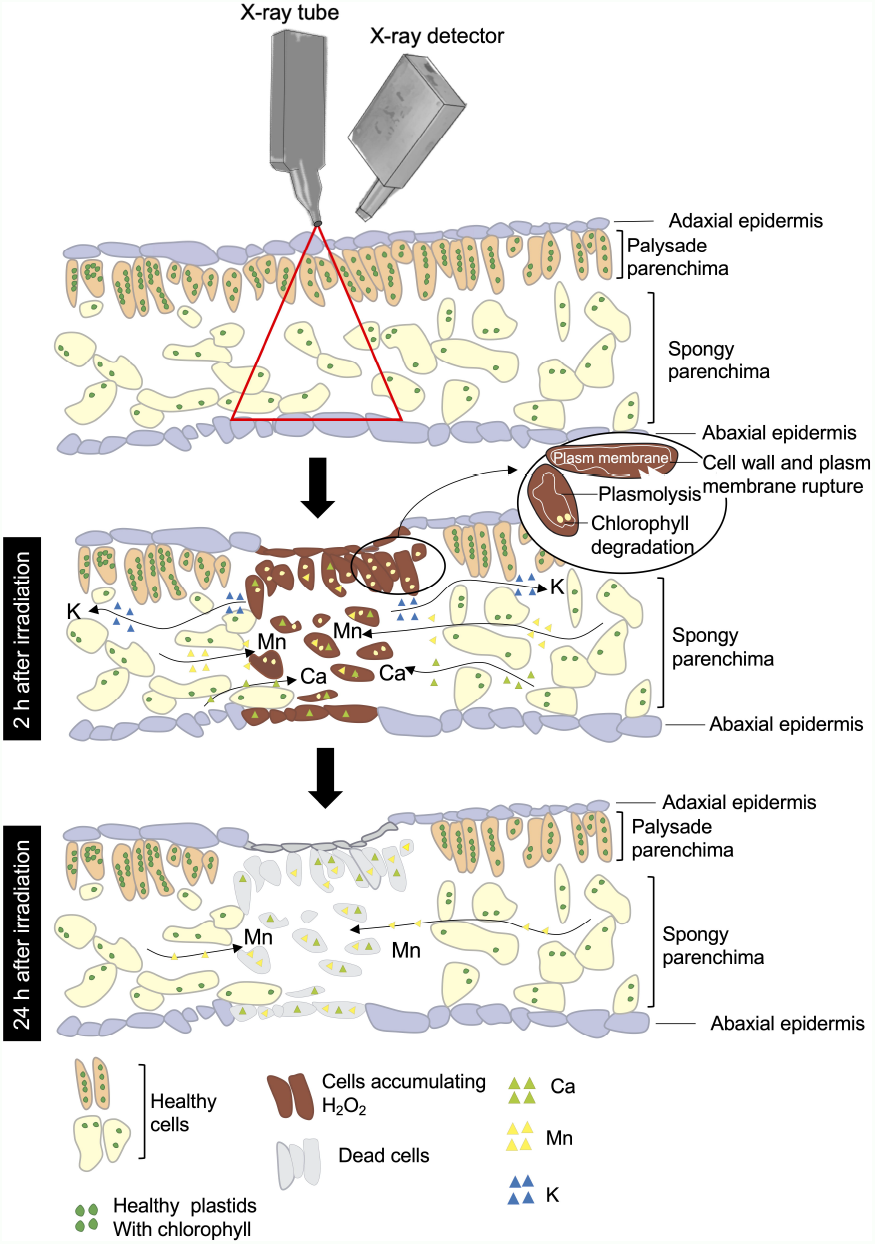
Time-resolved effects of high-doses X-ray exposure on soybean leaf tissues. During the first 2 hours after the exposure, the X-ray irradiation induces chlorophyll degradation, plasmolysis, and rupture of the cell walls and plasm membrane, thereby leading to K efflux, ROS accumulation and Ca and Mn increasing in the irradiated cells. However, most of the irradiated cells die 24 hours after exposure.

Finally, the characterisation herein presented reveals that a significant fraction of the elemental changes as a function of X-ray beam exposure stem from plants’ physiological signalling or response, and not simply from sample dehydration, as previously suggested. These response mechanisms are illustrated in the schematic representation shown in Fig. 6. Nevertheless, one should keep in mind that these effects are dose-dependent, and therefore, all studies employing X-ray or other radiation-based methodologies for biological samples are encouraged to include a preliminary investigation to verify if beam damage is interfering with the reliability of the data, and, if necessary, take the appropriate decision to overcome radiation artefacts.

## Supporting information

Electronic Supplementary Material

## SUPPLEMENTARY DATA

Supplementary data consist of the following. Fig. S1: setup employed for the *in vivo* XRF-based X-ray exposure assays on soybean leaves. Fig. S2: normalized XRF K, Ca, and Rh Lα count rate recorded on soybean leaves at V3 growth stage as a function of the exposure time (54 s, 24-60 min) a dose to a collimated 1 mm or polycapillary focused 30 µm X-ray beam derived from an Rh anode at 40.5 W (45 kV and 900 µA) or 4.5 power (45 kV and 100 µA), without or with a 25-µm thick Ti primary filter selected. Fig S3: normalized XRF K, Ca, Mn, and Rh Lα count rate recorded on soybean leaves at V3 growth stage exposed to polycapillary focused 30 µm X-ray beam at 40.5 W (45 kV and 900 µA) during two cycles of 20 minutes each, with a 2 h non-irradiation gap. Fig. S4: photographs and XRF probing of K, Ca, and Mn spatial distribution on fresh and dried soybean leaves at V3 growth stage exposed to polycapillary focused 30 µm X-ray beam at 40.5 W (45 kV and 900 µA) for 20 minutes. Fig. S5: XRF probing maps of K, Ca, and Mn spatial distribution on fresh and dried soybean leaves at the V3 growth stage from two independent biological replicates exposed to a polycapillary focused 30 µm X-ray beam at 40.5 W (45 kV and 900 µA) for 10 s. Fig. S6: XRF spectra recorded with and without the 25 µm-thick Ti primary filter selected on a plexiglass cube.

## ACKNOWLEDGMENTS

Dr. Helio Tolentino and Dr. Anna Paula Sotero, from the Brazilian Light Source Laboratory, are acknowledged for their invaluable assistance during the beam flux determination. Besides, we thank the Laboratory of Electron Microscopy “Prof. Elliot Watanabe Kitajima” at the Luiz de Queiroz College of Agriculture of the University of São Paulo for providing the infrastructure for the microscopy analysis, as well as Dr Eduardo de Almeida for his comments during the conduction of this work.

## AUTHOR CONTRIBUTIONS

**Conceptualization:** Montanha, GS and Carvalho, HWP; **Methodology:** Montanha, GS, Marques, JPR, Jones, MWM, and Carvalho; **Investigation:** Montanha, GS, Marques, JPR, and Rodrigues, ES; **Formal Analysis:** Montanha, GS, Marques, JPR, Jones, MWM, and Carvalho, HWP; **Data Curation:** Montanha, GS; **Resources:** Carvalho, HWP; **Funding Acquisition:** Carvalho, HWP; **Writing – Original Draft Preparation:** Montanha, GS; **Writing – Review & Editing:** Montanha, GS, Rodrigues, ES, Marques, JPR, Jones, MWM, and Carvalho;

## DATA AVAILABILITY

The raw data herein presented is fully available at the Figshare repository: https://doi.org/10.6084/m9.figshare.18584384.v1

## Abbreviations

[XRF]: X-ray fluorescence spectroscopy
[µ-XRF]: microprobe X-ray fluorescence spectroscopy
[TEM]: transmission electron microscopy
[DAB]: diaminobenzidine staining solution
[MDC]: monododecyl cadaverine staining solution
[ROS]: reactive oxygen species

