## Supplementary material for "Physiological responses of plants to *in vivo* XRF radiation damage: insights from anatomical, elemental, histochemical, and ultrastructural analyses": Electronic Supplementary Material

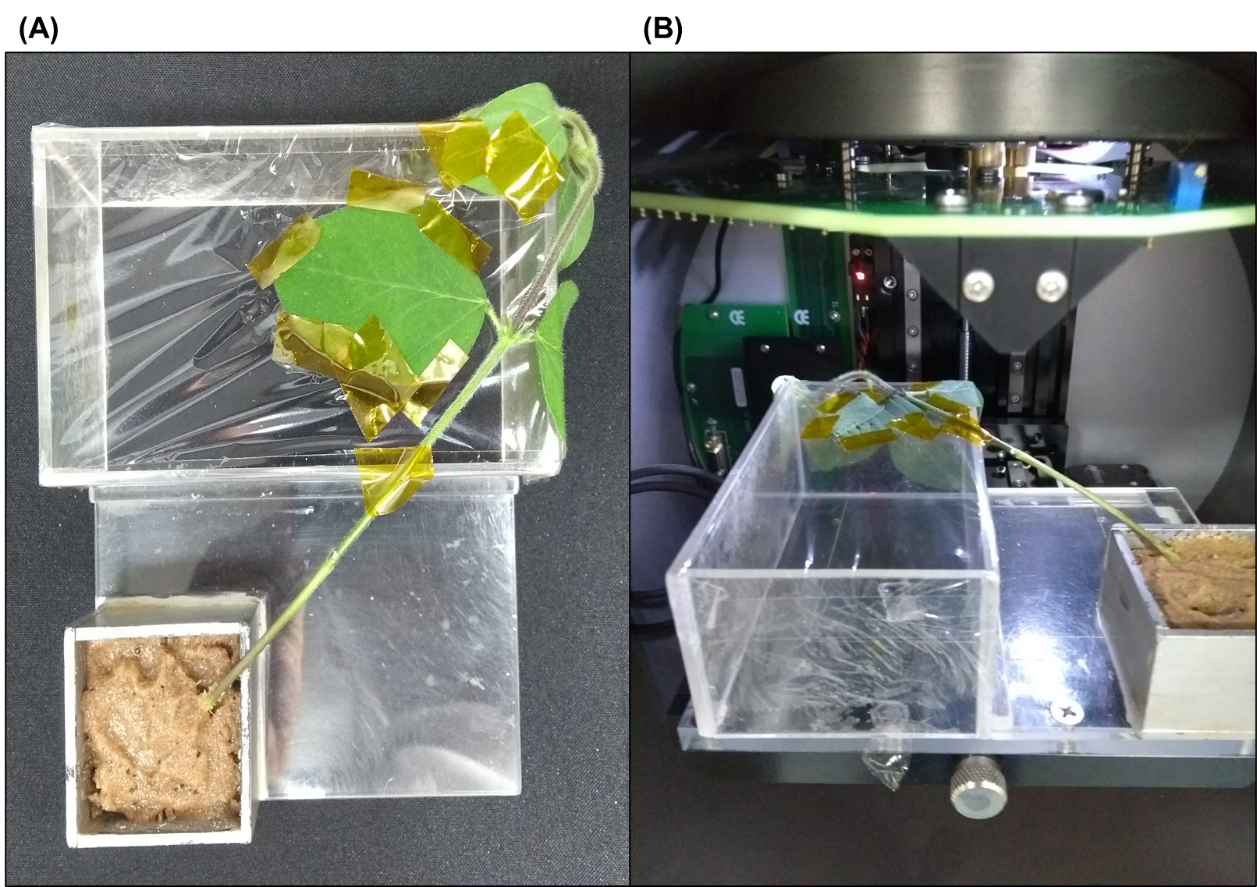

Figure S1. Setup employed for the *in vivo* XRF-based X-ray exposure assays on soybean leaves.

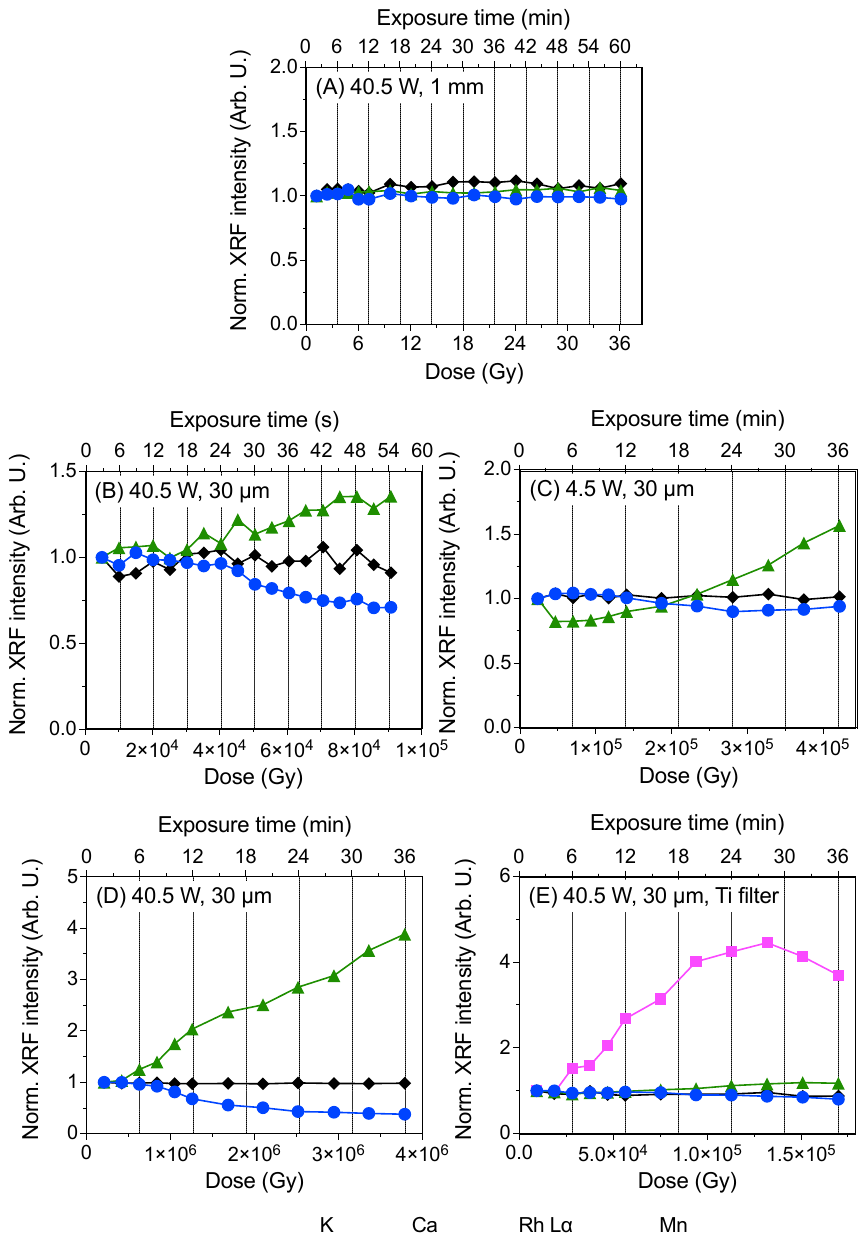

Figure S2. Normalized XRF K, Ca, and Rh Lα count rate recorded on soybean leaves at the V3 growth stage as a function of the time and radiation dose during the exposure either to a collimated 1 mm or polycapillary focused 30 µm X-ray beam derived from an Rh anode at 40.5 W (45 kV and 900 µA, A-C) or 4.5 power (45 kV and 100 µA, D), without (A-D) or with a 25-µm thick Ti primary filter selected (E). The count rates were normalized by their respectively first recorded values. Data resulting from an independent biological replicate.

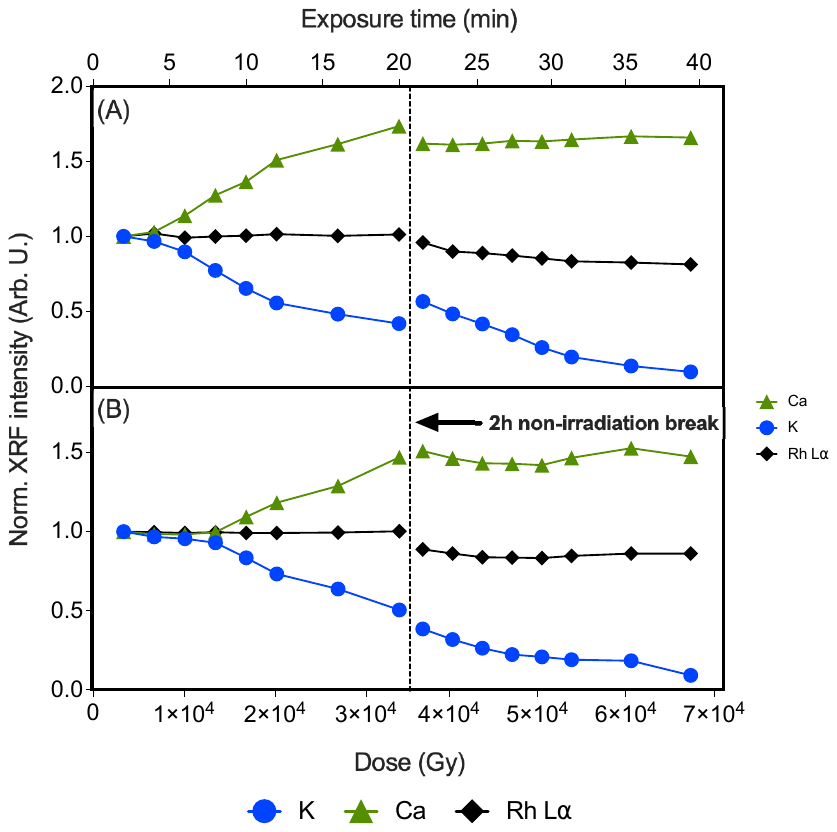

Figure S3. Normalized XRF K, Ca, Mn, and Rh Lα count rate recorded on soybean leaves at V3 growth stage exposed to polycapillary focused 30 µm X-ray beam at 40.5 W (45 kV and 900 µA) during two cycles of 20 minutes each, with a 2h non-irradiation gap. The figures (A-B) encompass independent biological replicates.

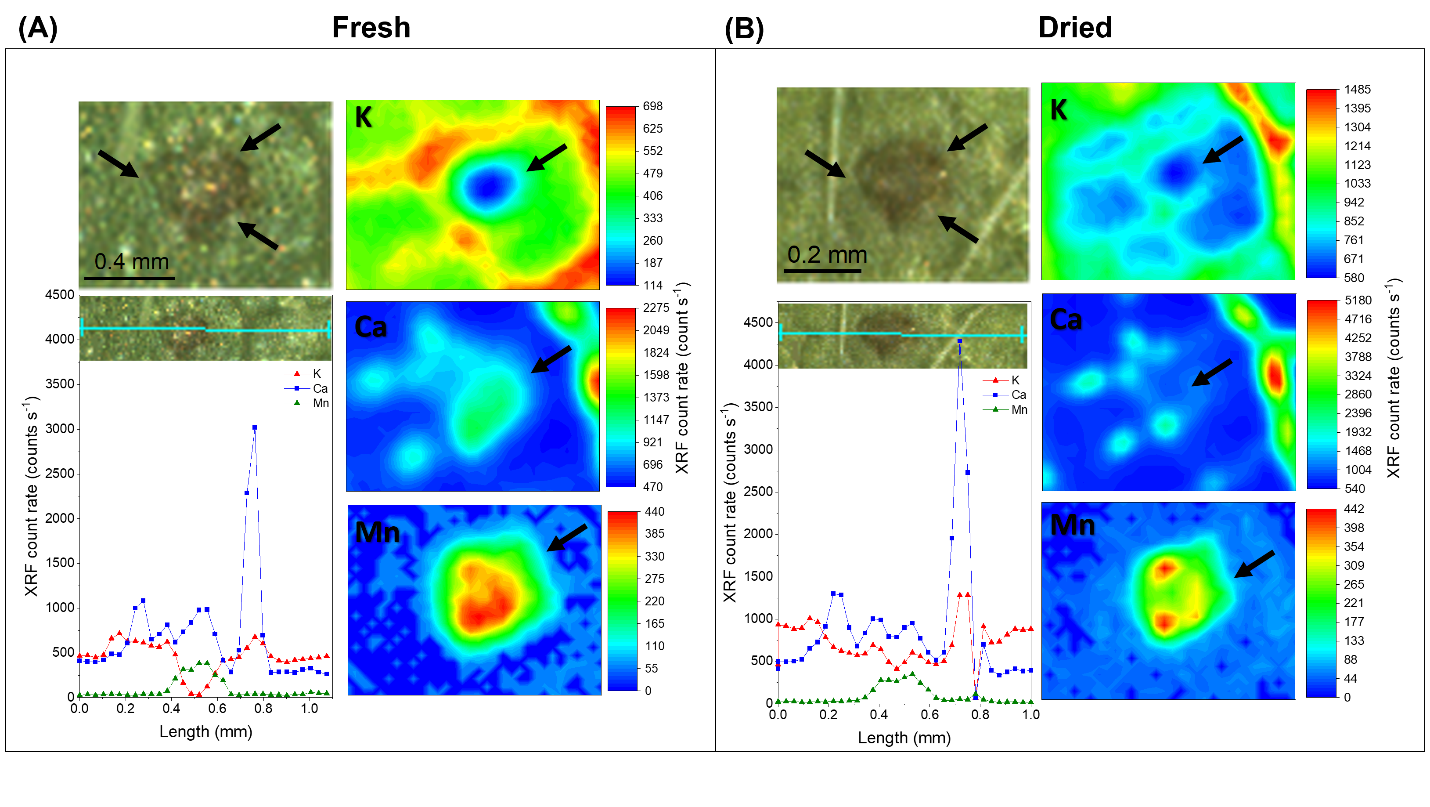

Figure S4. Photographs and XRF probing of K, Ca, and Mn spatial distribution on fresh (A) and dried (B) soybean leaves at V3 growth stage exposed to polycapillary focused 30 µm X-ray beam at 40.5 W (45 kV and 900 µA) for 20 minutes. An 800-pixel XRF map and a 32-point XRF linescan were carried out in each case. The analysis of the fresh leaf was carried out 24-h after the irradiations, whereas the dried ones after a 48-h oven-drying at 60 ºC. Note that the irradiated leaf regions, pointer out with the black arrows, presents a necrotic spot and clear changes on the elemental distribution in both fresh and dried leaf tissues. Data resulting from an independent biological replicate.

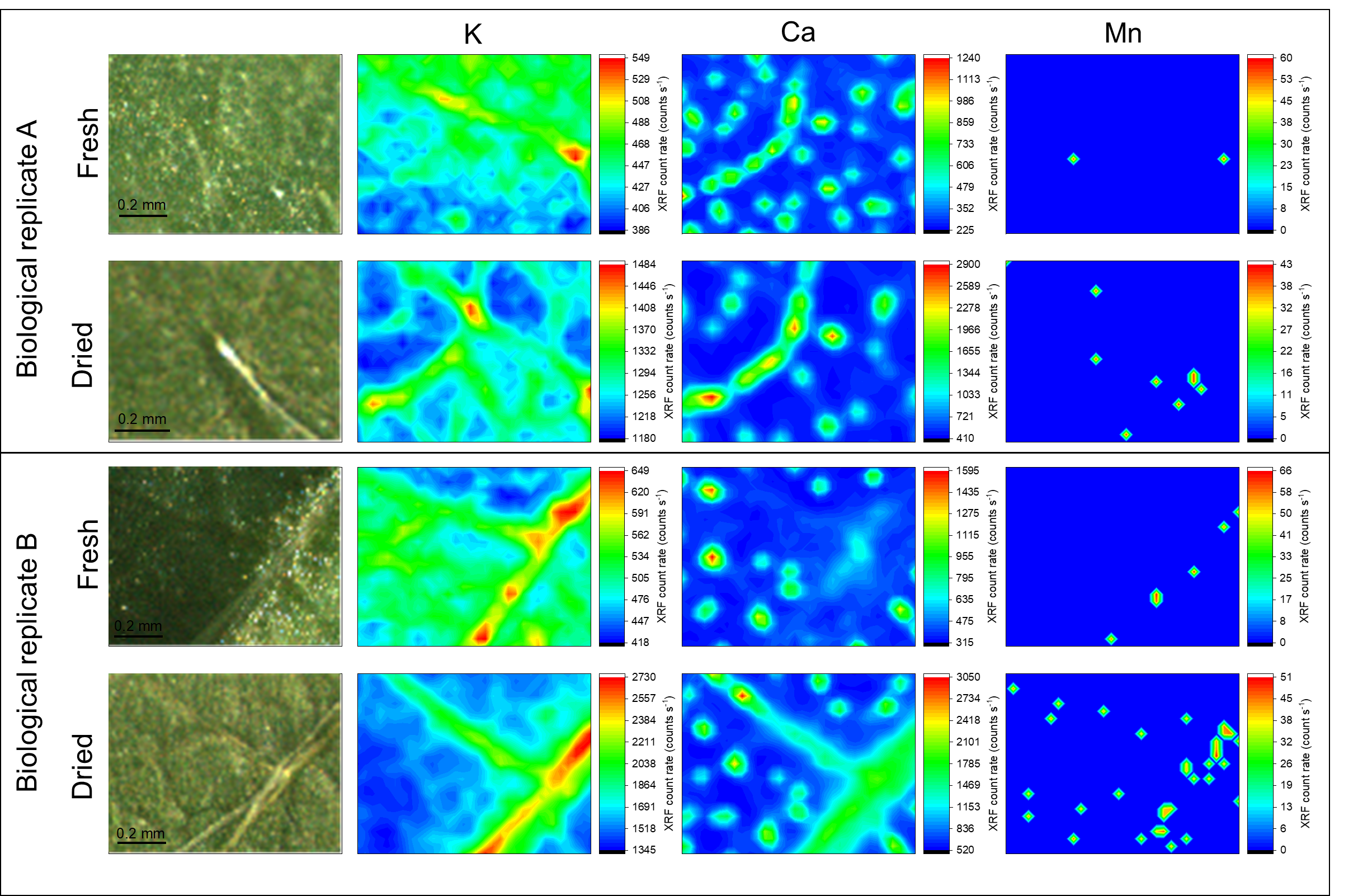

Figure S5. XRF probing maps of K, Ca, and Mn spatial distribution on fresh and dried soybean leaves at the V3 growth stage from two independent biological replicates exposed to a polycapillary focused 30 µm X-ray beam at 40.5 W (45 kV and 900 µA) for 10 s. The analysis of the fresh leaf was carried out 24-h after the irradiations, whereas the dried ones after a 48-h oven-drying at 60 ºC.

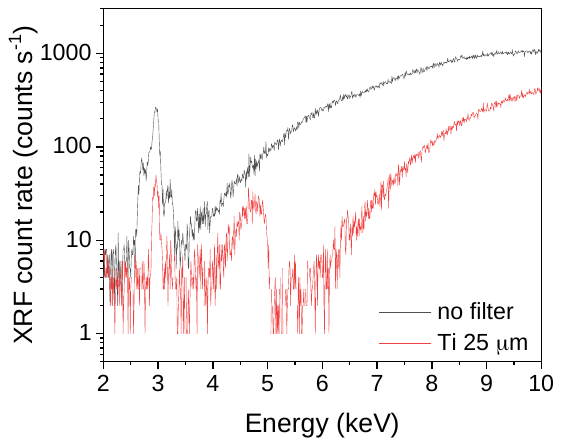

Figure S6. XRF spectra recorded with and without the 25 µm-thick Ti primary filter selected on a plexiglass cube. The use of filter implies on a lower flux, thus a decrease in X-ray scattering is noticed.
